# Beta-hydroxybutyrate promotes basal insulin secretion while decreasing glucagon secretion in mouse and human islets

**DOI:** 10.1101/2024.02.27.582117

**Authors:** Risha Banerjee, Ying Zhu, George P. Brownrigg, Renata Moravcova, Jason C. Rogalski, Leonard J. Foster, James D. Johnson, Jelena Kolic

**Affiliations:** Department of Cellular and Physiological Sciences, Life Sciences Institute, University of British Columbia Vancouver, Canada; Department of Biochemistry and Molecular Biology, University of British Columbia Vancouver, Canada

**Keywords:** Beta-hydroxybutyrate, beta cell, insulin, glucagon, human islets

## Abstract

Dietary carbohydrates raise blood glucose and limiting carbohydrate intake improves glycemia in patients with type 2 diabetes. Low carbohydrate intake (< 25 g) allows the body to utilize fat as its primary fuel. As a consequence of increased fatty acid oxidation, the liver produces ketones to serve as an alternative energy source. β-Hydroxybutyrate (βHB) is the most abundant ketone. While βHB has a wide range of functions outside of the pancreas, its direct effects on islet cell function remain understudied. We examined human islet secretory response to acute racemic βHB treatment and observed increased insulin secretion at low glucose concentrations (3 mM glucose). Because βHB is a chiral molecule, existing as both R and S forms, we further studied insulin and glucagon secretion following acute treatment with individual βHB enantiomers in human and C57BL6/J mouse islets. We found that acute treatment with R-βHB increased insulin secretion and decreased glucagon secretion at physiological glucose concentrations in both human and mouse islets. Proteomic analysis of human islets treated with R-βHB over 72 h showed altered abundance of proteins that may promote islet cell health and survival. Collectively, our data show that physiological concentrations of βHB influence hormone secretion and signaling within pancreatic islets.

## INTRODUCTION

The secretion of insulin and glucagon from the endocrine pancreas is essential for maintaining glucose homeostasis and regulating metabolism (1). Although glucose is the primary regulator of islet hormone secretion, other nutrients and metabolites may also play important roles (1). Although previous studies have investigated the effects of non-glucose substrates on islet cell function (2,3), some metabolites, such as ketone bodies, remain understudied.

Ketone bodies are produced as a byproduct of protein and fatty acid metabolism in the liver and are used as an energy source in the absence of glucose (4). Of the 3 ketone bodies (beta-hydroxybutyrate, acetoacetate, and acetyl coA) found in mammals, beta-hydroxybutyrate (βHB) is the most abundant (5). Racemic βHB is an equal mixture of its two naturally-occurring enantiomeric forms – R-βHB and S-βHB (5). R-βHB is the physiological enantiomer, while S-βHB, is isomerized in the liver into the physiological R enantiomer before being metabolized (5–7). The serum concentration of βHB in humans is <1 mM (6,8,9), but this level increases to > 2 mM during exercise, prolonged fasting and/or strict carbohydrate restriction, as seen with ketogenic diets (6, 10,11). This physiological state of ketosis is different from diabetic ketoacidosis (DKA), a serious medical condition where ketone concentrations in the blood can reach as high as 20 mM (12,13).

Investigations into the role of βHB in various cell types have shown that its metabolism in cardiomyocytes is upregulated for use as an energy source in the failing heart (14), and that βHB protects hepatocytes from ER-stress-induced cell death (15). Studies highlighting the benefits of the ketogenic diet have even led to an increase in the popularity of exogenous ketone supplements in an effort to further promote weight loss and slow the onset of type 2 diabetes (T2D) in prediabetic patients (16). Recently, clinical studies have shown that while exogenous ketone supplements lower blood glucose levels in metabolically healthy individuals (17), this effect does not extend to adults with T2D, where exogenous ketone supplements did not improve glycemic control or lower blood glucose following acute or chronic treatment (18).

However, the direct effect of βHB on pancreatic islet function has not been studied in detail. Early studies suggest that βHB can potentiate insulin release in INS1 cells (19) and rat islets in combination with glucose or other metabolites such as monomethyl succinate (20–23), while a recent study shows that ketone esters can stimulate insulin release in obese, but not lean, C57BL/6J mice (24). In study that employed human islets from 6 donors, βHB was shown to augment insulin release in the presence of 5.6 mM glucose (25). However, there is currently little to no evidence outlining the mechanism of effect of βHB on both insulin and glucagon secretion from human islets.

In the present study, we investigated the effect of acute and chronic βHB treatment on insulin and glucagon secretion in both human and mouse pancreatic islets. We found that acute R-βHB exposure increases insulin secretion and decreases glucagon secretion under physiological glucose concentrations in both human and mouse islets. Proteomic analysis revealed differential abundance of proteins involved in cell proliferation, amino acid biosynthesis and metabolism upon R-βHB treatment. The results of our study now implicate a role for βHB in altering hormone release in human and mouse islets. Considering the growing popularity of exogenous ketone supplements and the ketogenic diet, our study illustrates the need for additional investigations into the effects of ketone bodies on islet function and glucose homeostasis.

## MATERIALS AND METHODS

### Human islet culture

Human islets for research were provided by the Alberta Diabetes Institute IsletCore at the University of Alberta in Edmonton (www.bcell.org/adi-isletcore), used with approval of the Human Research Ethics Board at the University of Alberta (Pro00013094; Pro00001754) and the University of British Columbia (UBC) Clinical Research Ethics Board (H13-01865). All donors’ families gave informed consent for the use of pancreatic tissue in research. Information about individual donors can be found in Supplementary Table 1. Details regarding the islet isolation procedures can be found in the in the protocols.io repositor (26).

Each islet preparation was received in a 50 ml conical centrifuge tube containing CMRL culture media (Corning, 15-110-CV) (26). Upon arrival, islets were further purified by hand-picking under a stereomicroscope. Islets were then suspended in RPMI 1640 medium (Thermo Fisher Scientific, Cat#, 11879-020) supplemented with 5.5 mM glucose, 10% fetal bovine serum (FBS), and 100 units/mL penicillin/streptomycin, and cultured in 10 cm non-tissue culture coated (NTCC) petri dishes (ThermoFisher Scientific, Cat # FB0875713) overnight at 37°C, 5% CO_2_.

For static secretion experiments with racemic βHB, 260 islets were picked into NTCC dishes with 10 mL culture media. For static secretion experiments with individual enantiomers of βHB, 3 islets in 100 µL of culture media were picked into a 96 V-well plate (Corning: #CLS3894) per replicate. Islets were given ∼48 hours to adhere to the V-well prior to start of experiment.

### Mouse islet isolation and culture

C57BL/6J mice were purchased from the Jackson Laboratory and housed in the (temperature-controlled) UBC Modified Barrier Facility using UBC Animal Care Committee-approved protocols and following international guidelines. The mice were on a 12 h light/dark cycle and fed a chow diet (LabDiet #5053) with water *ad libitum*. Mouse islets were isolated by ductal collagenase (Type XI, Sigma, Cat. # C7657) injection as described previously (27). Islets were then hand-picked to purity. Mouse islets were cultured overnight in RPMI media with 11.1 mM D-glucose supplemented with 10% vol/vol fetal bovine serum (Thermo Fisher Scientific, Cat. #12483020) and 1% vol/vol Penicillin-Streptomycin (Gibco, Cat #15140-148) at 37°C with 5% CO_2_. For static secretion experiments, 3 islets in 100 µL of culture media were picked into a 96 V-well plate (Corning: #CLS3894) per replicate.

### Glucose stimulated insulin secretion of human islets treated with racemic βHB

Krebs-Ringer HEPES (KRBH) Buffer (pH 7.2, 129 mM NaCl, 4.8 mM KCl, 1.2 mM MgSO_4_, 1.2 mM KH_2_PO_4_, 2.5 mM CaCl_2_, 5 mM NaHCO_3_, 10 mM HEPES, 0.50% BSA, and ddH_2_O to volume) was prepared at glucose concentrations of 3 mM and 15 mM. 3 mM βHB was prepared from a 300 mM stock solution of β-hydroxybutyric acid sodium salt (Sigma-Aldrich, H6501) dissolved in water. Islets were then transferred into a new NTCC dish containing 10 mL 3 mM glucose KRBH and incubated at 37°C, 5% CO_2_ for 1 h. Following this, 12 samples of 20 islets were set up in 1.5 mL microcentrifuge tubes with 500 µL of 3 mM racemic βHB in either 3 mM glucose KRBH, 15 mM glucose KRBH, or the corresponding vehicle controls (each in triplicates), and incubated at 37°C, 5% CO_2_ for 30 min. The tubes were centrifuged at 1000 RPM for 1 min and the supernatant was collected and stored at −20°C; 500 µL of acid-ethanol extraction buffer (75 % of 95 % ethanol, 23.5 % concentrated acetic acid, 1.5 % concentrated HCl) was added to the cell pellet and stored at −20°C prior to being assayed for insulin using human insulin specific RIA (Millipore Cat# HI-14K).

### Glucose stimulated insulin secretion (GSIS) and glucagon secretion in human and mouse islets treated with individual enantiomers of βHB

KRBH Buffer (see above) was prepared at glucose concentrations of 3 mM, 6 mM, 10 mM, and 15 mM. 3 mM βHB was prepared from a 300 mM stock solution of R-β-Hydroxybutyric acid sodium salt (Santa Cruz Biotechnology Sigma-Aldrich, Cat# sc-229050) or S-3-Hydroxybutyric acid sodium salt (Santa Cruz Biotechnology Cat# sc-236887). Islets were treated with βHB in an acute or chronic manner. For chronic treatment, 3 islets/well in 96 V-well plates (Corning: #CLS3894) were incubated in either vehicle, 3 mM R-βHB, or 3 mM S-βHB for 72 h in RPMI media supplemented with 10% FBS and 5% penicillin/streptomycin at 37°C, 5% CO_2_.

For acute treatments, islets were plated as described above, but βHB was only present during the different glucose stimulations. Three technical replicates (islets from the same donor or mouse) were assayed for each treatment condition. Islets were treated with either enantiomer of βHB or vehicle in different glucose concentrations for 45 min. The supernatant was collected, and islets were stored in 100 µL of extraction buffer (75 % of 95 % ethanol, 23.5 % concentrated acetic acid, 1.5 % concentrated HCl) at −20 °C.

To measure glucagon secretion, 3 islets/well in 96 V-well plates (Corning: #CLS3894) were pre-incubated in 10 mM glucose for 1 h. Three technical replicates (islets from the same donor or mouse) were assayed for each treatment condition. Islets were treated with either enantiomer of βHB or vehicle in different glucose concentrations for 45 min. The supernatant was collected, and islets were stored in 100 µL of extraction buffer and 5 µg/mL of aprotinin (Sigma-Aldrich: A6279) at −20°C.

Mouse insulin was quantified using the rat insulin radioimmunoassay kit (Millipore Cat# RI-13K, RRID:AB_2884035). Human insulin was quantified using the human insulin radioimmunoassay (Millipore Cat# HI-14K, RRID:AB_2801577) or the human insulin ELISA kit (Crystal Chem Cat# 90095). Glucagon was quantified using radioimmunoassay (Millipore-Sigma, GL-32K, RRID:AB_2757819) or ELISA (Crystal Chem Cat#81520, RRID:AB_2884901) according to manufacturer’s instructions.

### Proteomics

Proteomic analysis was conducted according to established proteomics of the LSI Proteomics Core facility. Briefly, 3 mM solutions of either R-βHB or vehicle (sterile ddH_2_O) were prepared in human islet culture media and added to 6 cm petri-dishes (Falcon, Cat. # 351007). 200 islets were added to each plate, and incubated for 72 h at 37°C, 5% CO_2_. The islets were then picked into 1.5 mL microcentrifuge tubes and washed with 200 µL of 1X PBS. The islet pellets from each treatment were flash-frozen and stored at −70°C. Protein was extracted by treating frozen pellets with 60 µL of SDS lysis buffer (4 % SDS, 100 mM Tris, pH=8), vortexing, and then heating for 10 min at 100°C. the lysates were then centrifuged at 4°C and 10000 rpm for 10 min. Protein concentration was measured from the supernatant using the Pierce^TM^ BCA Protein Assay Kit (ThermoFisher Scientific, Cat # 23225). Islet lysate samples were run on a 10% SDS-PAGE gel, visualized using Coomassie Blue, and the bands were excised from the gel. The resulting proteins were reduced, alkylated, and digested by incubating with trypsin. The purified peptides were analyzed using a Bruker TIMS-ToF Pro II mass spectrometer coupled to a Bruker Nanoelute UPLC. The TIMS-ToF Pro II was set to Data-Independent Acquisition Parallel Accumulation-Serial Fragmentation (diaPASEF) scan mode for DIA acquisition scanning 100-1700 m/z. The capillary voltage was set to 1800V, drying gas to 3L/min, and drying temperature to 180°C. Acquired diaPASEF data were then searched using FragPipe computational platform (v. 17.1) with MSFragger (v. 3.4) (28), Philosopher (v. 3.8) (29), EasyPQP (v. 0.1.27) and DIA-NN (v. 1.8) to obtain DIA quantification, with use of the spectral library generated from a highly fractionated sample. Quantification mode was set to “Any LC (high precision).” All other settings were left default.

### Cell death screening

Mouse islets were isolated and dispersed into 384 well plates in culture media (11.1 mM D-glucose RPMI (Gibco, Cat#11875093), 1% vol/vol Penicillin/Streptomycin (GIBCO: 15140-148), 10% vol/vol FBS (Thermo: 12483020)) at a density of 5000 cells/well as previously described (27). Islet viability was measured with the TC20 Automated Cell Counter (Bio-Rad: 1450102) with Trypan Blue (Bio-Rad: 1450021). Cells were stained with 50 ng/mL Hoechst 33342 (Invitrogen) (to stain all cells), and 0.5 μg/mL propidium iodide (Sigma-Aldrich) in RPMI 1640 medium (Invitrogen) (to stain for dead cells) with 100 U/mL penicillin, 100 μg/mL streptomycin (Invitrogen), 10% vol/vol FBS (Invitrogen). After one hour of staining, the cells were treated with either vehicle, 3 mM R-βHB, or 3 mM S-βHB. The 384-well plate was placed in the environmentally controlled (37°C, 5% CO_2_) ImageExpress^MICRO^ high content imaging system. To measure cell death, the cells were imaged every 2-6 hrs for 96 h. Dispersed islet cells from C57BL6/J mice treated with 10 µM Thapsigargin in DMSO along with R-βHB, S-βHB, or vehicle (DMSO) were imaged every 2 h for 72 h. Cell death was quantified as the number of propidium iodide positive cells/number of Hoescht 33342 positive cells.

### Statistics and data analysis

GraphPad Prism 9 was used to conduct statistical analyses. Grubb’s test (a=0.05) was used to identify and eliminate statistical outliers. Insulin and glucagon secretion data were analyzed using a simple linear regression model on GraphPad Prism. Significance was assessed by Kruskal-Wallis test and Dunn’s multiple comparison test at 95% confidence, using p<0.05 to define statistical significance.

Differences between treatment groups were compared using 2-tailed t-test of log_2_(abundance). Only proteins detected in >80% of donors were considered for analysis. A p-value cutoff was applied to identify significance in differential abundance of proteins between treatments (p<0.05, fold change ≥1.2 for upregulated proteins, and fold change ≤0.8 for downregulated proteins) (Supplementary table 2). The 15 proteins displaying the highest or lowest fold-changes and p<0.05 between treatment conditions were uploaded to WebGestalt (30), a KEGG-based gene ontology enrichment software, to identify biological processes involving these proteins.

## RESULTS

### βHB increases basal insulin secretion in human islets

The intention of this study was to investigate the effects of acute and chronic βHB treatment on insulin and glucagon secretion in both human and mouse pancreatic islets (Figure 1A). We first performed a concentration-response pilot study, where we examined insulin secretion in islets obtained from cadaveric human donors (n=3, ADI IsletCore, UAlberta). Racemic βHB started to affect insulin secretion at 3 mM, as evidenced by a decreased stimulation index (**Supplementary Figure 1**). We next examined insulin secretion in response to acute 3 mM racemic βHB treatment at both basal (3 mM) and stimulatory (15 mM) glucose concentrations in human islet donors. We observed a 44% decrease in fold-insulin secretion (n=22, p<0.01, **Figure 1B**), however this was strictly driven by an increase in insulin secretion at low glucose (**Figure 1C**), since insulin secretion at 15 mM glucose was unchanged (**Figure 1D**). Total insulin content remained unchanged upon βHB treatment (**Figure 1E**).

**Figure 1.**
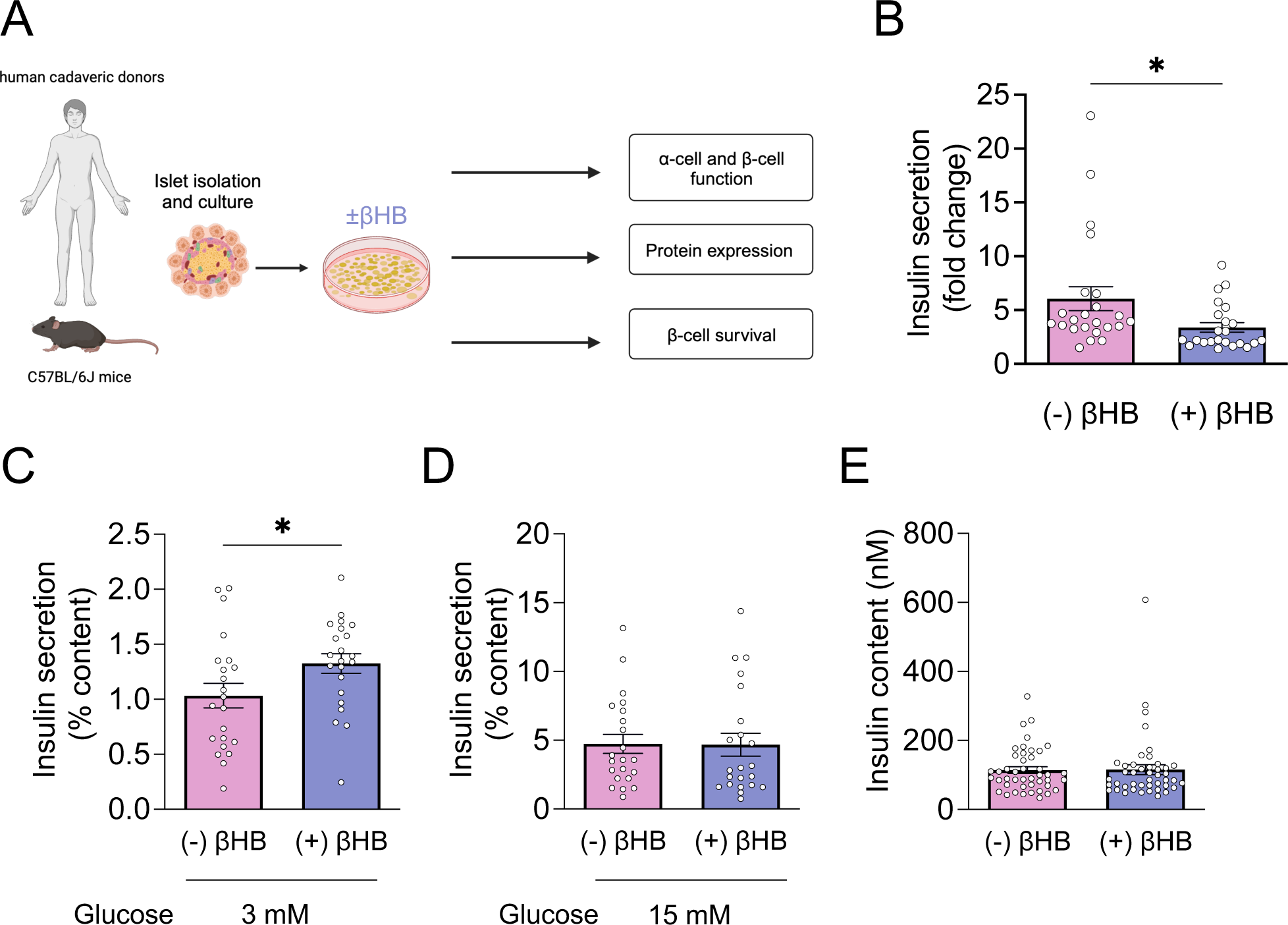
Experimental overview and effect of βHB on insulin secretion in human islets. **(A)** Experimental design for study of βHB on high-quality primary human islets isolated from cadaveric donors and from C57BL6/J mice. **(B)** Overall fold-change insulin secretion (15 mM / 3 mM glucose) in islets from cadaveric human donors (n = 22) with or without 3 mM racemic βHB. **(C)** Insulin secretion in islets from cadaveric human donors (n = 22) treated acutely with 3 mM racemic βHB at low (3 mM) glucose. **(D)** Insulin secretion in islets from cadaveric human donors (n = 22) treated acutely with 3 mM racemic βHB at high (15 mM) glucose. **(E)** Total **i**nsulin content from islets used in (B) is shown. Bars represent mean ± SEM. Each circle represents the mean of three technical replicates (Technical replicates refer to multiple islets from the same donor). Significance levels are indicated as asterisks over the corresponding bars. * indicates p<0.05.

Considering that R-βHB is the physiological enantiomer in humans (5), we investigated whether βHB affected hormone secretion from human islets in an enantiomer-specific manner. To also study whether the effects of βHB enantiomers are glucose concentration-dependent, islets from cadaveric human donors (n=10, ADI IsletCore, UAlberta) were treated acutely with either 3 mM R-βHB or S-βHB at multiple glucose concentrations (3, 6, 10, and 15 mM). To account for heterogeneity in individual islet size and composition, we normalized insulin secretion to the total islet insulin content, which was statistically unchanged overall following our acute treatments (**Supplementary Figure 2**). We observed a 1.7-fold increase in insulin secretion in islets treated with 3 mM R-βHB under basal glucose conditions (3 mM) (n=10, p<0.01, **Figure 2A**). We did not observe a change in insulin secretion upon 3 mM S-βHB treatment, nor was there a change in insulin secretion upon treatment with either enantiomer of βHB at glucose concentrations at or above 6 mM glucose concentrations (**Figure 2A**). However, we did observe considerable inter-islet differences in response to βHB, consistent with our recent study showing considerable donor-donor heterogeneity in insulin secretion in response to macronutrient stimuli (2). To explore this further, we stratified insulin secretion in response to either R-βHB or S-βHB treatment by donor BMI. While there was also no significant change in mean insulin secretion or in βHB -potentiated insulin secretion at 3 mM glucose in response to 3 mM R- βHB (**Figure 2B**), we did observe a 5-fold decrease in ΒHB-potentiated insulin secretion (fold change compared to vehicle) in islets from donors with BMI >25 (overweight or obese) at a suprathreshold glucose concentration of 6 mM and R-βHB treatment (p<0.001, **Figure 2C**). In S-βHB treated islets, there was a ≥2-fold decrease in insulin secretion in islets from donors with BMI > 25 compared to those from donors with BMI < 25 at both 3- and 6-mM glucose (**Figure 2D, E**), suggesting an enantiomer-specific BMI-dependency in insulin secretion in human islets. We then repeated this experiment in islets from donors with T2D. While we observed trends similar to those observed in islets from donors without diabetes, the limited availability of donors with T2D, restricts these observations to islets from a single donor (**Supplementary Figure 3A**). Overall, our results indicate that while racemic βHB leads to an increase in insulin secretion in human islets, this increase is driven by R-βHB at physiological glucose concentrations and is influenced by individual donor characteristics such as BMI (**Supplementary Figure 4**).

**Figure 2.**
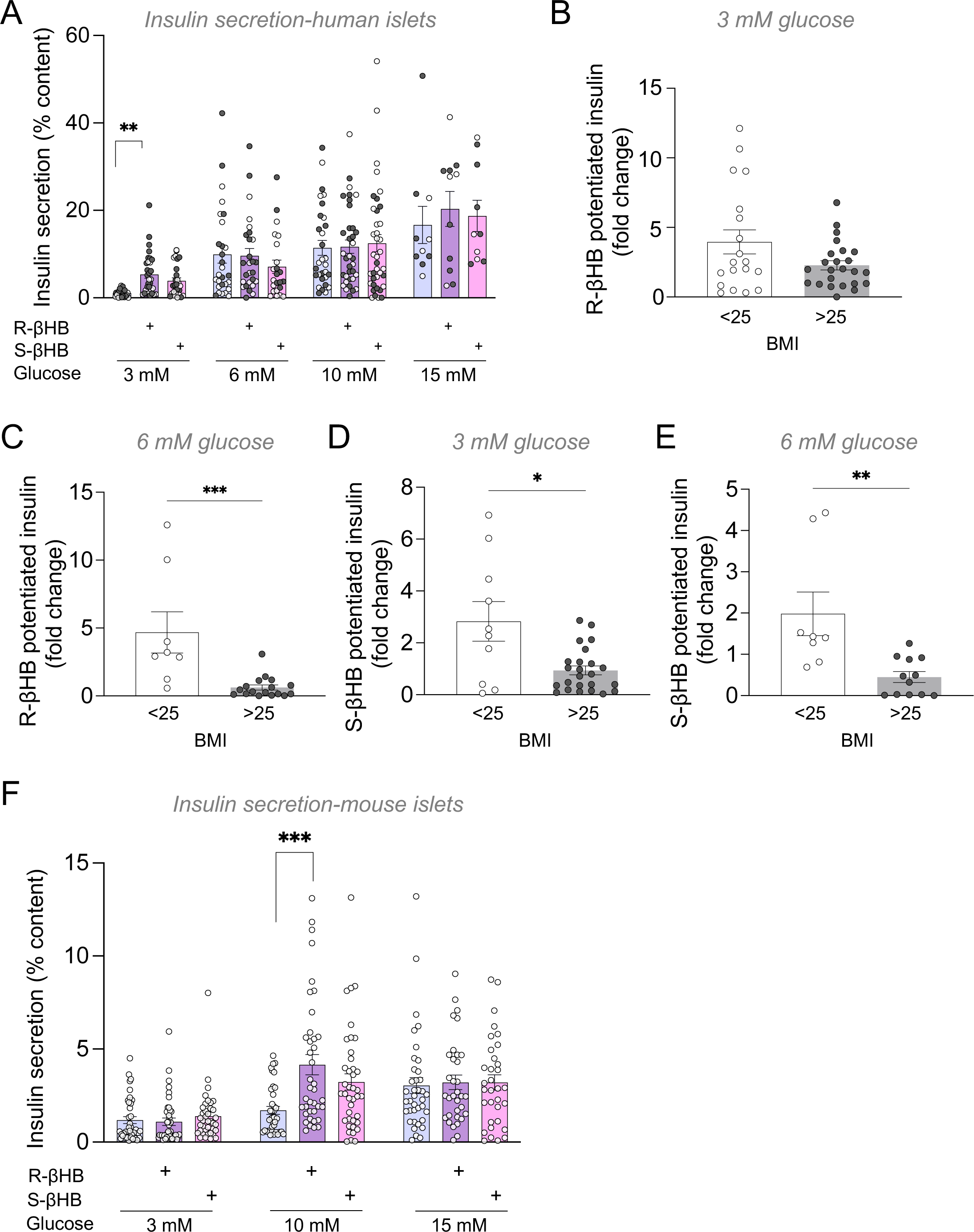
The effect of individual enantiomers of βHB on insulin secretion in human and mouse islets. **(A)** Insulin secretion as % of total islet content in islets from cadaveric human donors (n=10) treated acutely with 3 mM R- or S- βHB at 3 mM, 6 mM, 10 mM or 15 mM glucose is shown. Each circle represents a technical replicate. Black circles correspond to secretion values from donors with BMI > 25 and white circles correspond to secretion values from donors with BMI <25. **(B)** R-βHB-potentiated fold-change in insulin secretion values in islets treated with 3 mM glucose and 3 mM R-βHB is shown for donors with BMI > 25 or <25 from (A). **(C)** R-βHB-potentiated fold-change in insulin secretion values in islets treated with 6 mM glucose and 3 mM R-βHB is shown for donors with BMI > 25 or <25 from (A). **(D)** S-βHB-potentiated fold-change in insulin secretion values in islets treated with 3 mM glucose and 3 mM S-βHB is shown for donors with BMI > 25 or <25 from (A). **(E)** S-βHB-potentiated fold-change in insulin secretion values in islets treated with 6 mM glucose and 3 mM S-βHB is shown for donors with BMI > 25 or <25 from (A). **(F)** Insulin secretion as % of total islet content in islets from C57BL6/J mice (n=13) treated acutely with 3 mM R- or S- βHB at 3 mM, 10 mM or 15 mM glucose is shown. Each circle represents a technical replicate (Technical replicates refer to multiple islets from the same donor or same mouse). Significance levels are indicated as asterisks over the corresponding bars. * indicates p<0.05, ** indicates p<0.01, *** indicates p<0.001, and **** indicates p<0.0001.

### βHB increases glucose-stimulated insulin secretion in mouse islets

Given that previous studies have identified a role for βHB in potentiating insulin release in rodent islets (31), we next sought to investigate any enantiomer-specific effects of βHB on islets from C57BL/6J mice. Islets were treated acutely with either 3 mM R-βHB or S-βHB at glucose concentrations of 3 mM, 10 mM, or 15 mM. At 10 mM glucose, we observed a 2.6-fold increase in insulin released upon R-βHB treatment (p<0.01, n=13) (**Figure 2F**). Interestingly, there was no change in insulin release upon treatment with S-βHB at 10 mM glucose, or with either enantiomer at 3- or 15-mM glucose (**Figure 2F**), suggesting species-specific differences in ketone body-potentiated insulin secretion.

### βHB decreases glucagon secretion in both mouse and human islets

Glucagon secretion from islets is dependent on local glucose concentrations. In a hyperglycemic environment, insulin can suppress glucagon secretion via the PI3K/Akt signaling pathway (32). We thus next assessed glucagon secretion in human islets treated with 3 mM R- βHB, or S-βHB at glucose concentrations of 3 mM, 6 mM, 10 mM, and 15 mM. Interestingly, we observed a ∼46% decrease in glucagon secretion in islets treated with R-βHB, but not with S- βHB. (p<0.05, n=11, **Figure 3A**), implicating a direct role of ketone bodies on glucagon secretion. There was no change in glucagon secretion in islets treated with either enantiomer in the presence of higher glucose concentrations when we examined all donors; there was also no change in mean glucagon secretion in a BMI-dependent manner. However, when we again stratified βHB-mediated fold-changes in glucagon secretion by donor BMI, we observed a 1.5-fold increase in glucagon secretion at 10 mM glucose (**Figure 3B**). In islets treated with S-βHB, we also observed higher βHB-potentiated glucagon secretion (fold change compared to vehicle) at 10 mM glucose in donors with BMI > 25 compared to those with BMI > 25 (p<0.01, **Figure 3C**). Interestingly, in islets treated with S-βHB at 15 mM glucose, donors with BMI > 25 displayed lower βHB-potentiated glucagon secretion compared to those with BMI< 25 (p<0.01, **Figure 3D**). We repeated these experiments in islets from a single diabetic human donor (**Supplementary Figure 3B**) – while we again see trends in glucagon secretion similar to those seen in donors without diabetes, our observations are limited to n=1. When we examined islets isolated from C57BL6/J mice, we observed a 23% decrease in glucagon secretion upon treatment with R-βHB at 10 mM compared to vehicle control (n=13, p<0.01, **Figure 3E**). There was no change in glucagon release in islets treated with R-βHB at 3- or 15 mM glucose, or S-βHB at any glucose concentration (**Figure 3E**). Overall, our results indicate that R-βHB treatment leads to a decrease in glucagon secretion at physiological glucose concentrations in both human and mouse islets. Similar to our observations in the context of insulin secretion in human islets, this change might be influenced by donor characteristics such as BMI.

**Figure 3.**
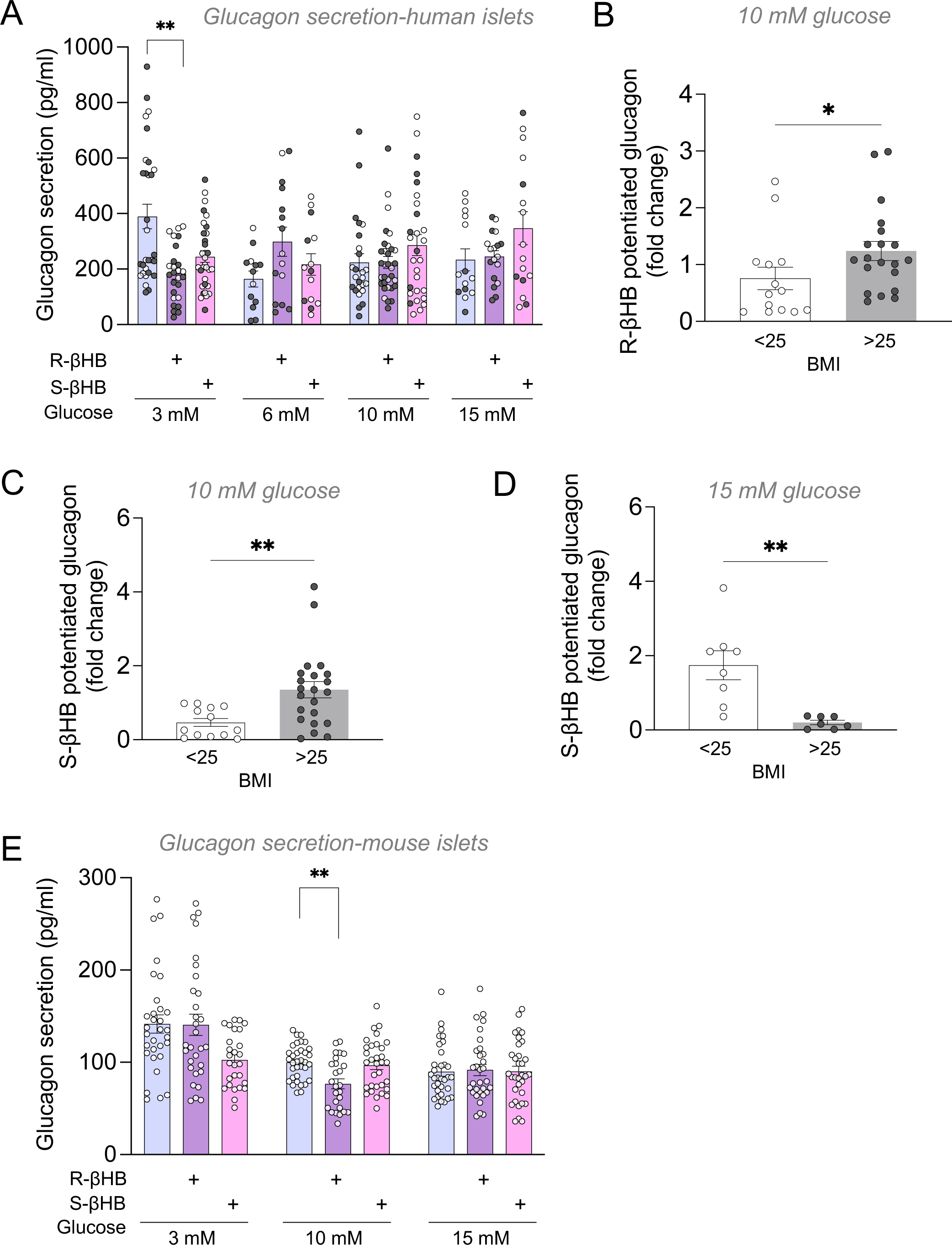
The effect of individual enantiomers of βHB on glucagon secretion in human and mouse islets. **(A)** Glucagon secretion in islets from cadaveric human donors (n=10) treated acutely with 3 mM R- or S- βHB at 3 mM, 6 mM, 10 mM or 15 mM glucose is shown. Each circle represents a technical replicate. Black circles correspond to secretion values from donors with BMI > 25 and white circles correspond to secretion values from donors with BMI <25. **(B)** R-βHB-potentiated fold-change in glucagon secretion in islets treated with 10 mM glucose and 3 mM R- βHB is shown for donors with BMI > 25 or <25 from (A). **(C)** S-βHB-potentiated fold-change in glucagon secretion in islets treated with 10 mM glucose and 3 mM S-βHB is shown for donors with BMI > 25 or <25 from (A). **(D)** S-βHB-potentiated fold-change in glucagon secretion in islets treated with 15 mM glucose and 3 mM S-βHB is shown for donors with BMI > 25 or <25 from (A). **(E)** Glucagon secretion in islets from C57BL6/J mice (n=9) treated acutely with 3 mM R- or S- βHB at 3 mM, 10 mM or 15 mM glucose is shown. Each circle represents a technical replicate (Technical replicates refer to multiple islets from the same donor or same mouse). Significance levels are indicated as asterisks over the corresponding bars. * indicates p<0.05, ** indicates p<0.01, *** indicates p<0.001, and **** indicates p<0.0001.

### Chronic βHB treatment does not affect insulin secretion in human islets

We next tested whether prolonged exposure to βHB impact hormone secretion from human islets. Islets were treated with either R- or S- βHB for 72 h prior to measuring glucose-stimulated insulin secretion or glucagon secretion at 3-, 6-, and 10-mM glucose. Neither enantiomer of βHB had a significant effect on insulin secretion or glucagon secretion at any of the glucose concentrations tested (**Supplementary Figure 5**). However, it is interesting to note that 4/5 donors we examined had a BMI over 25.

### βHB impacts cellular proliferation and nutrient-response signaling pathways

To characterize the molecular pathways affected by βHB treatment in human islets, we performed LC-MS-based proteomics. Human islets were treated with R-βHB for 72 h. We focused on R-βHB specifically because under chronic conditions we would expect S-ΒHB to be converted to R-βHB within the body (5). Islets were treated with R-βHB for 72 h to account for changes in protein abundance in cases where protein turnover time may not be limited to a few hours (33). Our proteomic analysis identified >7000 unique proteins, of which 201 displayed significant differences in abundance upon R-βHB treatment (p<0.05). Upon stratifying by fold change, 81 gene products were either upregulated (fold change > 1.2), or downregulated (fold change <0.8) following βHB treatment (**Figure 4A, B**). Web Gestalt Over-Representation Analysis (ORA) revealed that proteins involved in response to ketones (CDK4 (Cyclin-dependent kinase 4), FOXO1(Forkhead Box Protein O1), TXNIP (Thioredoxin Interacting protein), and hormone receptor binding (SOCS2 (Suppressor of Cytokine Signaling 2)) are enriched in islets treated with R-βHB. We also observed an enrichment in biological processes associated with cellular response to drugs (CDK4, FOXO1, SSH1 (Protein phosphatase Slingshot homolog 1)) and the Wnt signaling pathway ( FOXO1) (**Supplementary Figure 6**). TXNIP has also been shown to play a role in inducing apoptotic cell death β-cells, consequently leading to a decrease in insulin secretion (34). Additionally, DNAL1 (Dynein Axonemal Light Chain 1) and CEP131 (Centrosomal Protein of 131 kDa), primarily associated with β-cell ciliary movement in response to glucose (35), had lower abundances upon R- βHB treatment. Interestingly, the proteins FOXO1 and PDCD2 were increased in abundance in R-βHB-treated islets: these proteins are all involved in islet response to free fatty acids such as palmitate. In mice, FOXO1 is involved in maintaining metabolic homeostasis in the islet and regulates insulin signaling in response to oxidative stress, contributing to β-cell compensation (36). Collectively, these data identify numerous differentially abundant proteins upon chronic R-βHB treatment and implicate several cellular signaling pathways including regulation of cell death and inflammatory signaling, autophagy, vesicle transport, and fatty acid-induced signaling.

**Figure 4.**
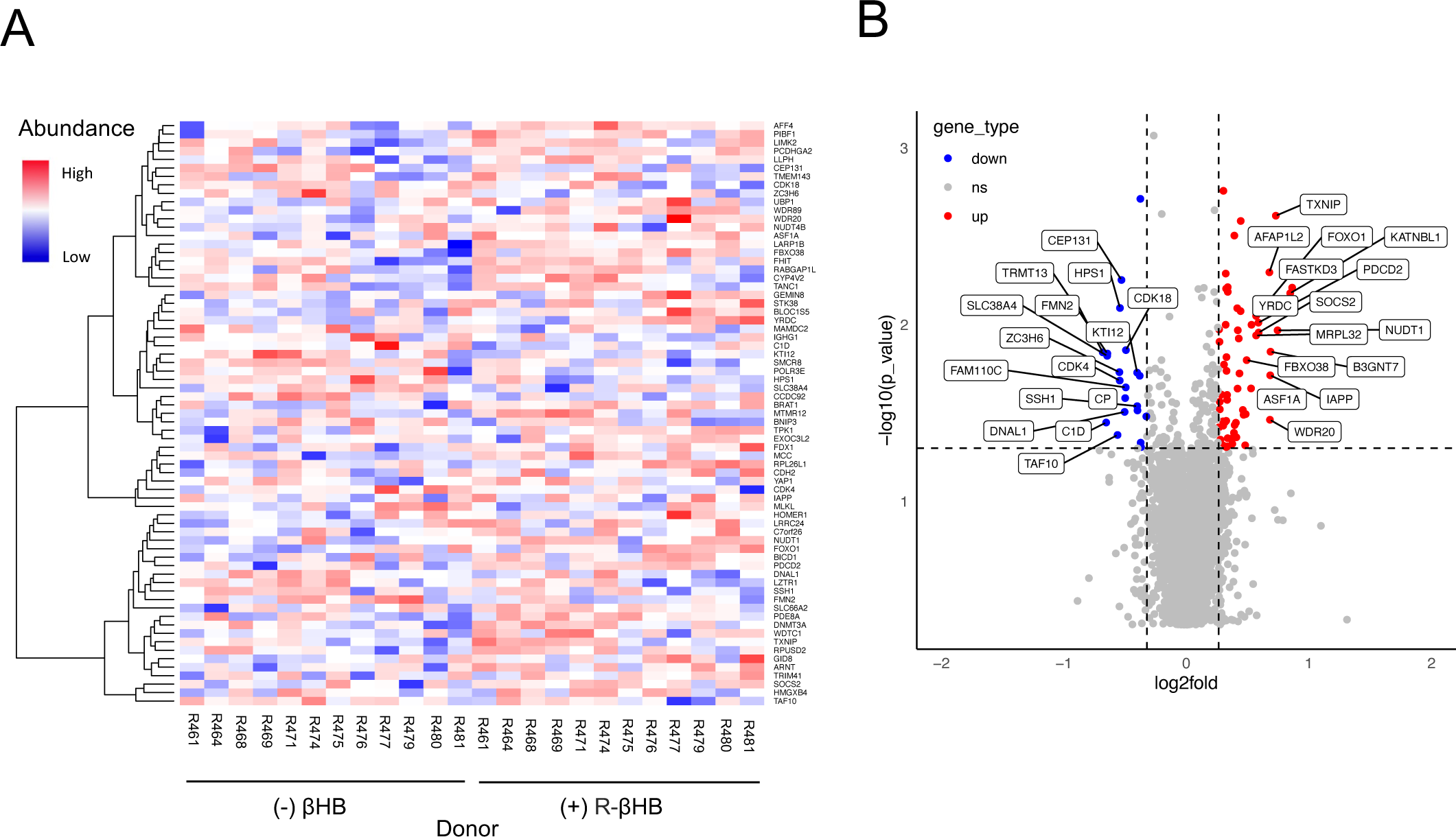
Proteomic analysis of human islets treated with R-βHB. **(A)** Heatmap showing fold change (vs. vehicle) of proteins displaying differential abundance (p<0.05) upon prolonged R- βHB treatment (72 h) in human islets (n=12 cadaveric human donors). **(B)** Volcano plot showing proteins displaying differential abundance upon chronic R-βHB treatment in human islets. Top 15 proteins displaying highest and lowest fold change are labelled-upregulated proteins are shown in red and downregulated proteins are shown in blue. Grey dots indicate proteins that do not show significant differential expression between treatment conditions (ns).

### Chronic βHB treatment does not protect islet cells from ER-stress induced cell death

The results of our proteomics analysis revealed a decreased abundance of proteins involved in cell death and inflammatory signaling. βHB has been shown to promote cell survival and proliferation in hepatocytes (15). Previous studies have also shown that chronic exposure to βHB attenuates the unfolded protein response (UPR) in rat liver cells and counteracts ER-stress-induced inflammatory cascades in hepatoma cells, thus improving cell viability (15,37). Thus, to determine whether either enantiomer of βHB has a similar protective effect on islet cells, we used kinetic cell death screening to measure cell death in response to prolonged R-βHB and S-βHB exposure. We observed no significant effect of treatment with either enantiomer of βHB compared to vehicle controls in islets from both male and female mice under the conditions we tested (**Figure 5A, B**, respectively). Islet cells from male mice had high % PI-positive cells after 96 h of treatment with either enantiomer of βHB; however, female mice did not exceed 40% cell death under the same conditions (**Supplementary Figure 7A, B**). We repeated this experiment using an ER stressor, thapsigargin (TG), in combination with βHB treatments to determine whether βHB can protect islet cells from ER stress-induced cell death. Dispersed islets from C57BL6/J mice (n=3, 2 male, 1 female mouse) were treated with either enantiomer of βHB in the presence and absence of 10µM TG. We observed no significant change in cell death in islet cells treated with either R or S-βHB compared to vehicle both in the presence and absence of TG overall (**Figure 5C, D**), and between individual male and female mice (**Supplementary Figure 7C, D**). Treatment with TG+βHB did not have a significant effect on islets from female mice, in contrast with islets from male mice, which showed a modest but significant reduction in cell death upon treatment with TG + R-βHB compared to TG + S-βHB, suggesting a sex-specific and enantiomer-specific effect of βHB on islet cell survival upon ER stress that requires further investigation (**Supplementary Figure 7E, F**). This agrees with previous studies, which have shown that female β-cells are more resilient to factors inducing cell-death, such as ER stress (27). These observations indicate that there may be slight differences in the effect of individual enantiomers of βHB on islet survival compared to each other, but that neither enantiomer plays significant roles in mouse islet cell survival, both in the presence or absence of the ER stress.

**Figure 5.**
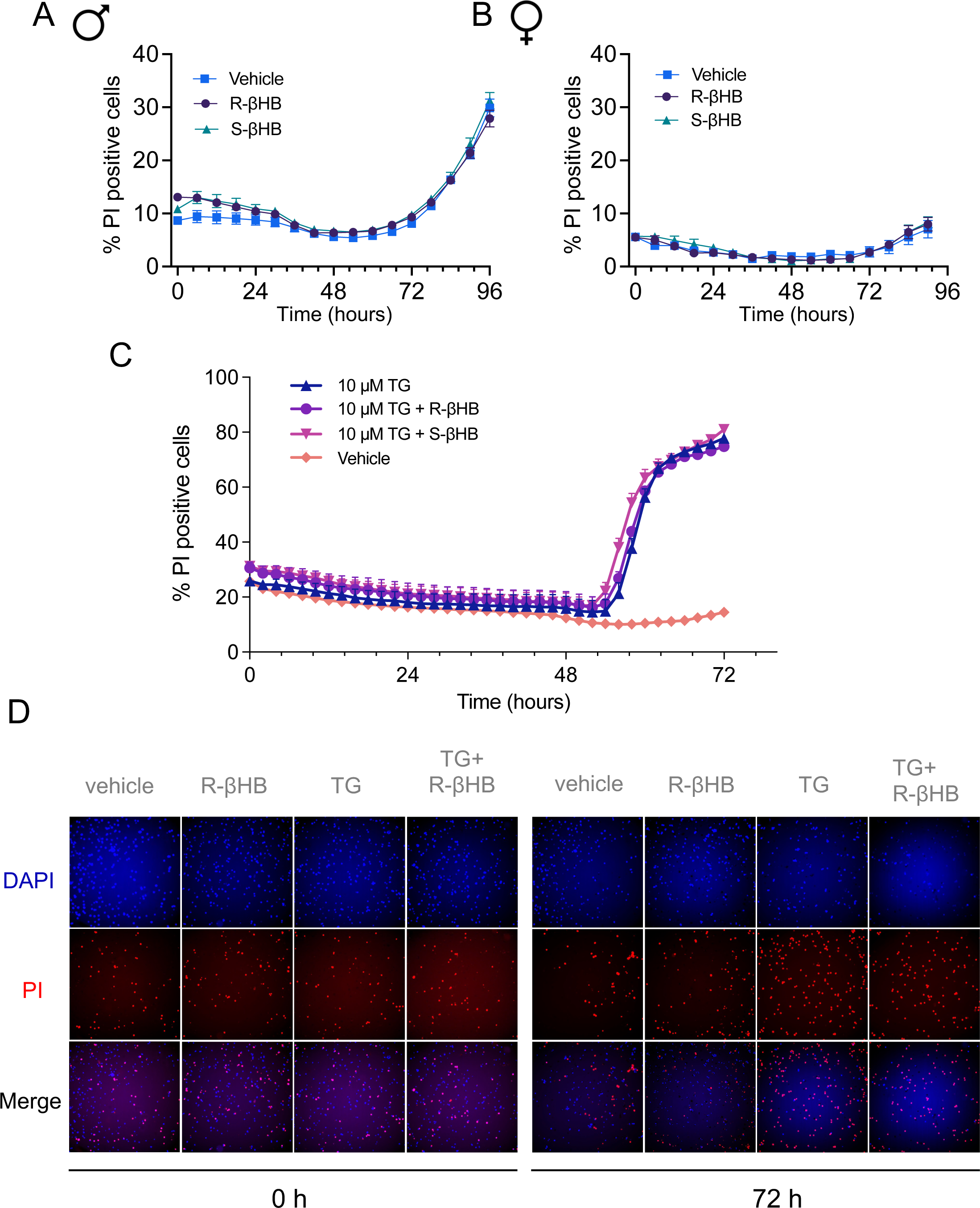
Effect of mouse islet cell survival following chronic βHB treatment. **(A, B)** Cell death as % PI positive cells in dispersed islet cells treated with either R- or S-βHB in male **(A)** or female **(B)** mice. **(C)** Cell death as % PI positive cells in dispersed islet cells treated with the ER stressor thapsigargin ± R-βHB or S-βHB is shown. **(D)** Representative images of dispersed islet cells at 0 h and 72 h. Results shown are quantified from n=6 male and n=2 female mice over 2 separate runs.

## DISCUSSION

The aim our study was to systematically investigate the effects of βHB on islet hormone secretion in both human and mouse pancreatic islets. In human islets, we found that acute treatment with racemic βHB increased insulin secretion at 3 mM glucose – this effect was primarily driven by R-βHB - the physiological enantiomer of βHB. R-βHB also led to a corresponding decrease in glucagon secretion under basal glucose conditions. Importantly, heterogeneity in donor characteristics influenced the effect of βHB on human islets; islets from donors with BMI greater than 25 (overweight/obese) had a smaller increase in insulin secretion compared to islets from donors with BMI less than 25. Interestingly, when we examined the effect of βHB under more chronic conditions (72 hr treatment), we did not see any significant effects on hormone secretion. However, the majority of the donors studied under 72-hr βHB treatment had elevated BMIs, which may have precluded the effects on glucagon and insulin secretion seen previously.

While the effects on islet hormone secretion seen in our study were subtle, several recent reports have suggested that elevated insulin levels under basal glucose conditions may be an early indicator of β-cell dysfunction and may contribute to the adaptive insulin resistance seen in individuals living with T2D (38–40). Moreover, both basal and nutrient-stimulated insulin secretion from human islet donors is highly variable (2). As such it is critical we understand how and if these subtle changes in islet hormone secretion upon βHB exposure, are physiologically relevant on an individual basis.

In mouse islets, R-βHB also potentiated insulin secretion and decreased glucagon secretion, however this was only seen at moderate glucose concentrations of 10 mM. S-βHB also slightly potentiated insulin secretion at 10 mM glucose, but this effect did not reach statistical significance. While we only studied islets from young, lean chow fed C57BL/6J mice, our results are consistent with a recent study that examined the effects of 10 mM R- and S-βHB on dynamic insulin secretion in isolated islets from both lean and obese C57BL/6J mice (24). This suggests that the effects of ketone bodies on islet hormone secretion are species-specific and further re-iterates the importance of studying human islets to improve our understanding of human islet biology.

Next, to discern molecular mechanisms underlying islet response to chronic βHB treatment and identify novel pathways, we performed proteomic analysis on human islets treated with R-βHB for 72 hours. Despite not seeing significant differences in islet hormone secretion following 72-hour exposure to βHB, we focused on a more chronic effect with our proteomics dataset to ensure that we captured any differences in protein abundances where protein turnover time may be longer than a few hours (33,41). Several proteins involved in cell death signaling and nutrient-response pathways were observed to have differential abundance in βHB-treated islets. As such, we too investigated whether either enantiomer of βHB would protect islet cells from (or contribute to) cell death under conditions of cellular stress. Because previous reports had suggested that that chronic exposure to βHB attenuates the unfolded protein response in other cell types (15,37), we focused on a model of endoplasmic reticulum stress. Surprisingly, our results showed that neither enantiomer of βHB appeared to protect islet cells from cell death. However, we did observe that female islets displayed a much lower cell death compared to their male counterparts, in support of previous studies on ER-stress mediated islet cell death (27). Our proteomics data indicate an increase in SOCS2 abundance, a suppressor of cytokine signalling, and a downregulation of CDK4, which is involved in cell cycle regulation at the G1/S checkpoint. Therefore, future experiments should investigate whether βHB treatment protects islet cells from cytokine-induced cell death.

While our results collectively show that that the effect of acute R-βHB on both human and mouse islet insulin and glucagon secretion is significant, donor BMI heavily influenced the strength of response. Chronic βHB treatment, indicative of either a prolonged ketogenic diet or chronic consumption of exogenous ketone supplements, altered the abundance of proteins involved in nutrient response and cell death in islets, however, neither enantiomer of βHB significantly improved islet cell survival. Considering the growing popularity of the ketogenic diet and availability of exogenous ketone esters, our findings now contribute to the mechanistic understanding of how increasing concentrations of ketone bodies may affect pancreatic islet cell signalling and function. In the face of chronic basal hyperinsulinemia being causal to numerous health conditions (46,47), we particularly hope that our results will encourage clinical researchers to look for individual response heterogeneity to increased plasma ketones.

## Author Contributions

R.B Conducted experiments, analyzed data, and wrote the manuscript.

Y.Z Conducted experiments and analyzed data

G.P.B conducted experiments and analyzed data (cell death screening)

R.M Conducted experiments and analyzed data (proteomics)

J.C.R Conducted experiments and analyzed data (proteomics)

L.J.F Supervised work, edited manuscript

J.D.J Conceived studies, supervised/guarantees the work, edited manuscript

J.K Conceived studies, conducted experiments, supervised/guarantees the work, edited manuscript.

## Acknowledgements

We thank members of the Johnson, Foster and MacDonald labs. We thank the Human Organ Procurement and Exchange (HOPE) program and Trillium Gift of Life Network (TGLN) for their work in procuring human donor pancreas for research, and James Lyon, Nancy Smith and Dr. Jocelyn Manning Fox (Alberta) for their efforts in human islet isolation. We especially thank the organ donors and their families for their kind gift in support of diabetes research.

