## Supplementary figures for "Beta-hydroxybutyrate promotes basal insulin secretion while decreasing glucagon secretion in mouse and human islets"

### Supplementary Figure 1

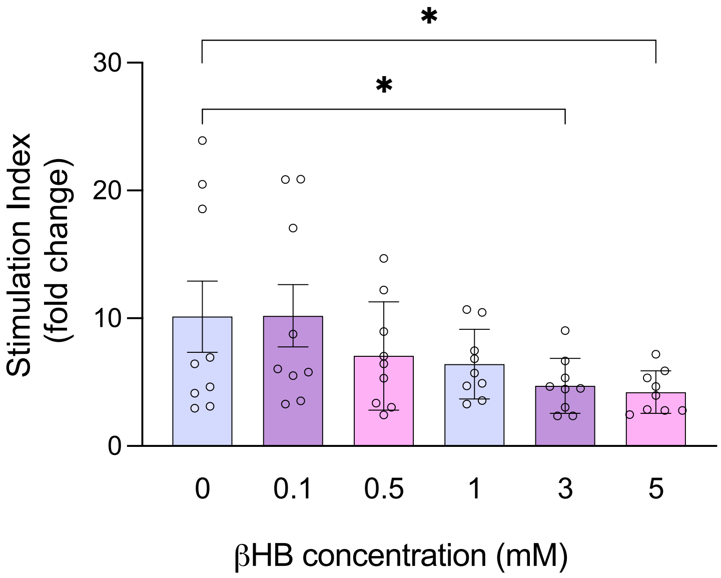


**Supplementary Figure 1**. **βHB Concentration-response curve.** Stimulation indices (high glucose/low glucose) in human islets (n=3 donors) treated with indicated concentrations of βHB. Each circle represents a technical replicate. (Technical replicates refer to different islets from the same donor). Significance levels are indicated as asterisks over the corresponding bars. * indicates $p<0.05$.

Supplementary Figure 2

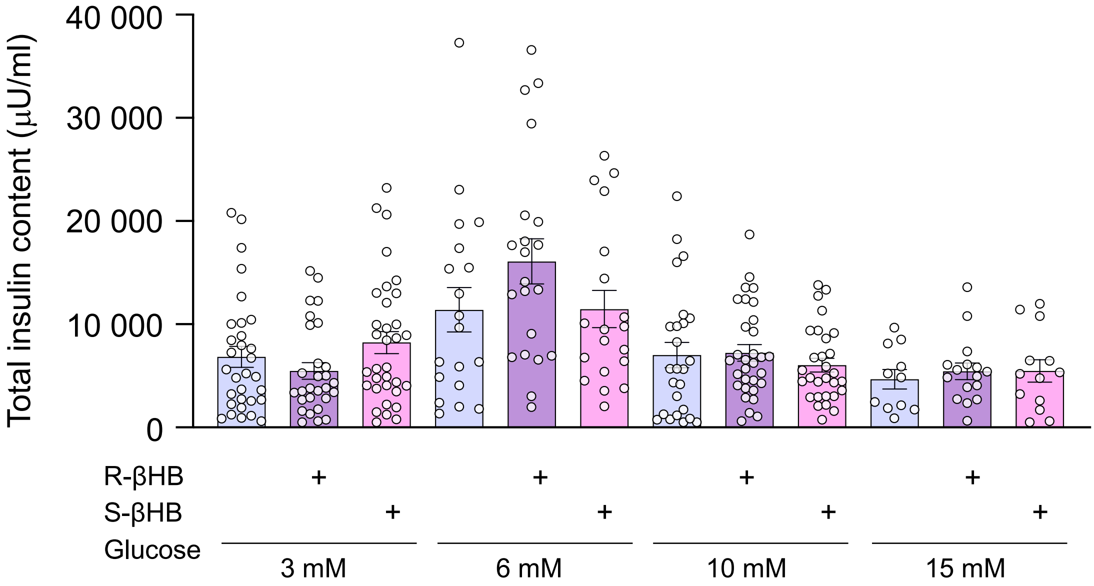


**Supplementary Figure 2**. **Total insulin content following acute R- or S-βHB treatment.** Insulin content from islets isolated from cadaveric human donors (n=10) treated with 3 mM R-βHB or S-βHB is shown. Each circle represents a technical replicate.

### Supplementary Figure 3

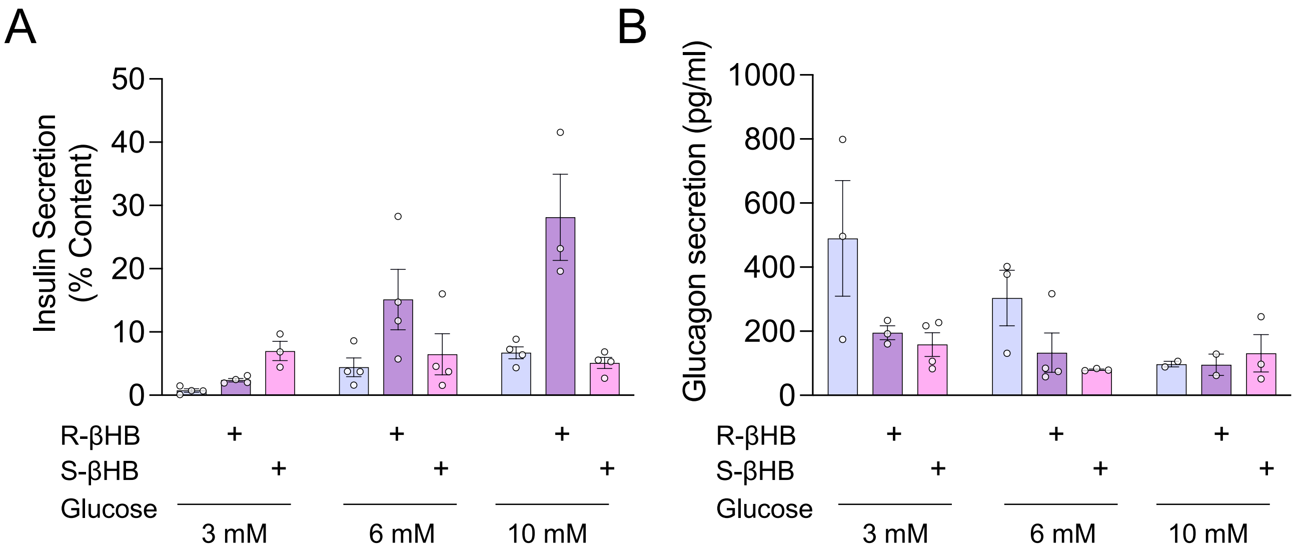


**Supplementary Figure 3. Effect of βHB on islet hormone secretion in T2D.** Insulin and glucagon secretion in islets treated with individual enantiomers of βHB in islets isolated from a single human donor with T2D (R452). **(A)** Insulin secretion as % of total islet content in islets from cadaveric human donors with T2D (n=1) treated acutely with 3 mM R- or S-βHB at 3 mM, 6 mM, and 10 mM glucose is shown. **(B)** Glucagon secretion in islets from cadaveric human donor with T2D (R452) (n=1) treated acutely with 3 mM R- or S-βHB at 3 mM, 6 mM, or 10 mM glucose is shown. Each circle represents a technical replicate.

### Supplementary Figure 4

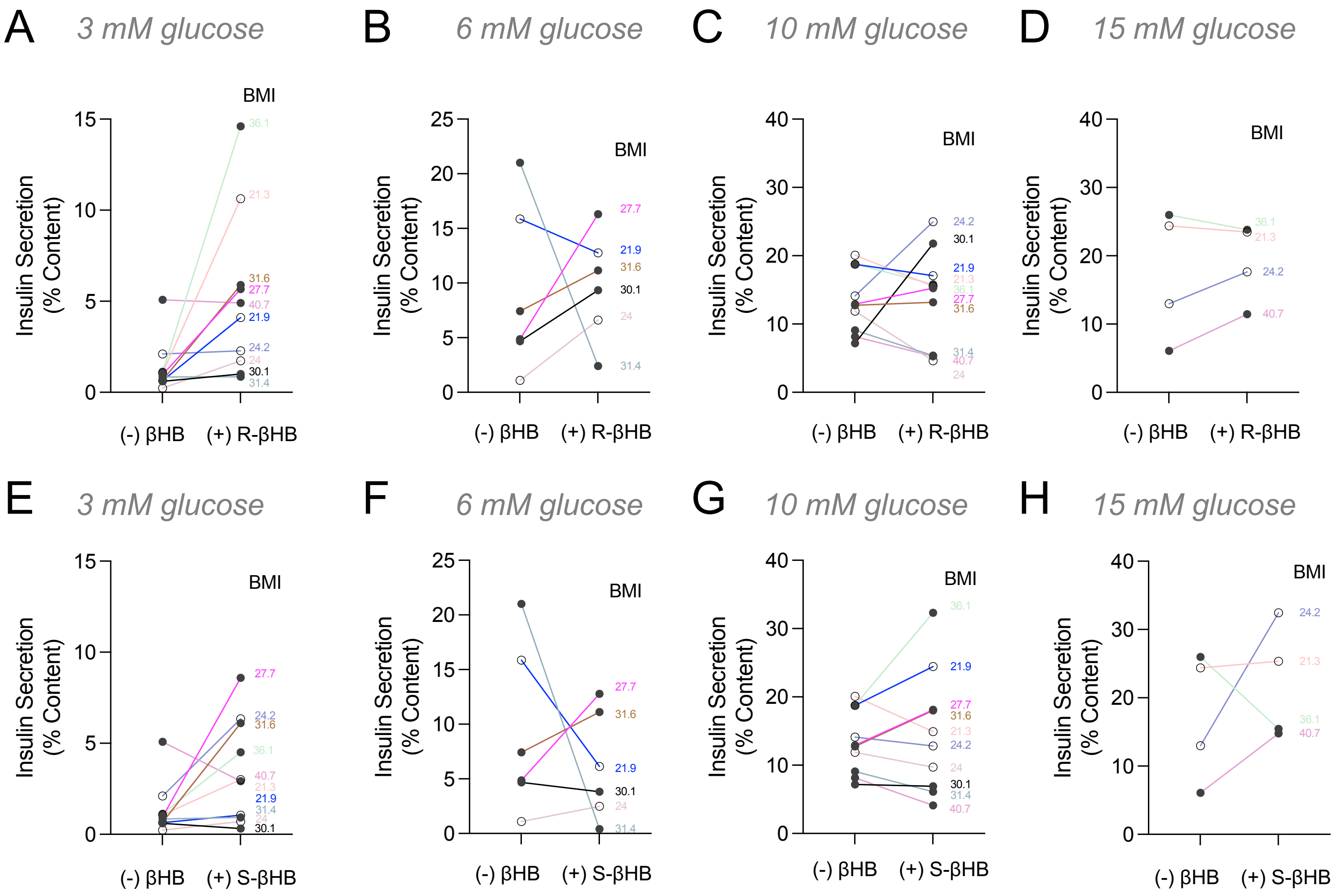


**Supplementary Figure 4. Effect of βHB on islet hormone secretion stratified by BMI.** Insulin secretion as % of total islet content in human islets treated acutely (45 min) with 3 mM R- or S-βHB, stratified according to donor BMI. Hollow circles indicate insulin secretion from donors with BMI <25, and full circles indicate insulin secretion from donors with BMI>25. **(A-D)** Insulin secretion in human islets treated with 3 mM R-βHB at 3 mM glucose (A), 6 mM glucose (B), 10 mM glucose (C), or 15 mM glucose (D). **(E-H)** Insulin secretion in human islets treated with 3 mM S-βHB at 3 mM glucose (E), 6 mM glucose (F), 10 mM glucose (G), or 15 mM glucose (H). Each colour corresponds to an individual donor. The BMI of individual donors is listed to the right of each plot. Each point is the average of 3-4 technical replicates.

### Supplementary Figure 5

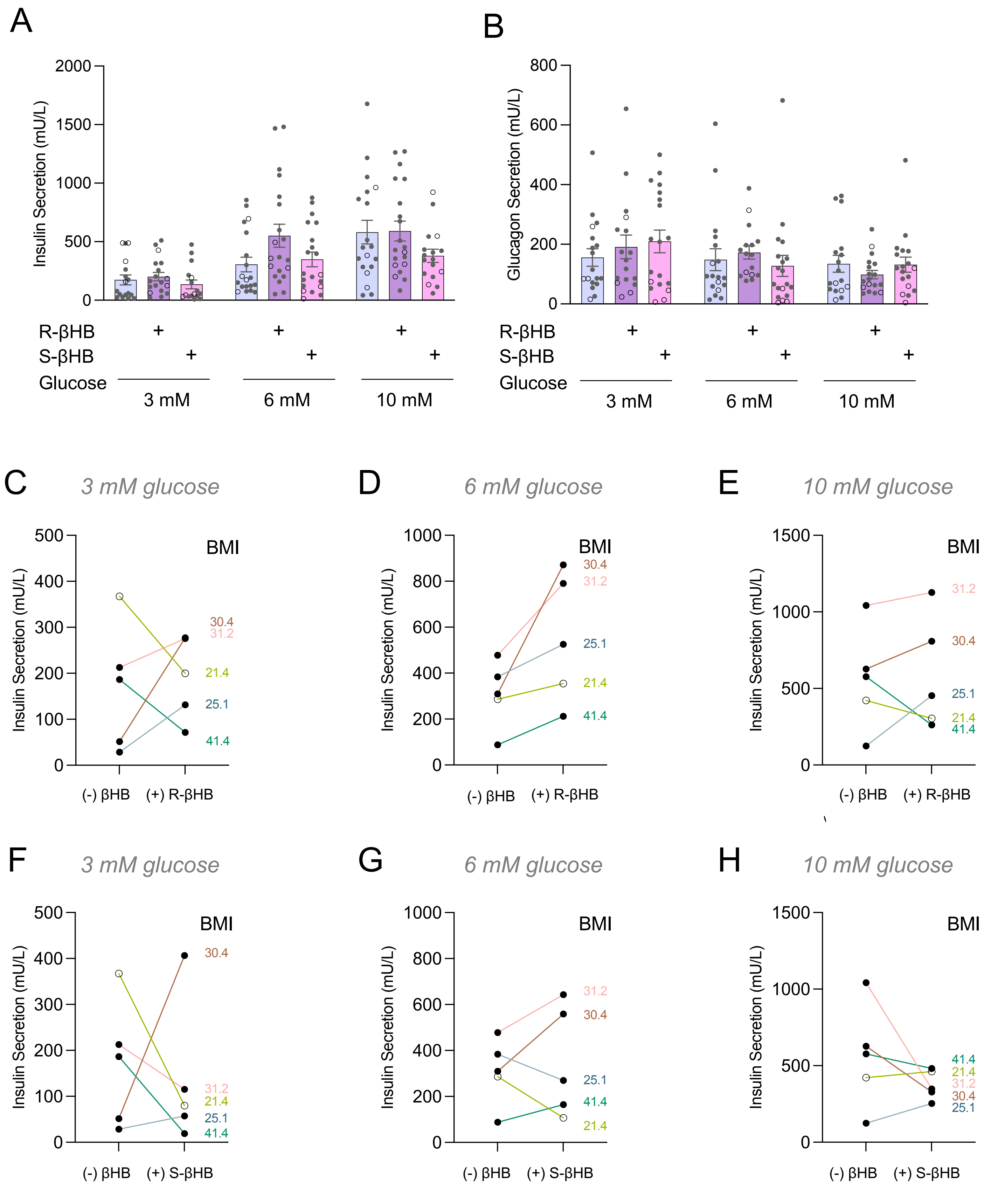

**Supplementary Figure 5**. **Hormone secretion in human islets following chronic exposure to βHB. (A)** Insulin secretion in human islets (n=5) treated with 3 mM R- or S-βHB for 72 h at 3-, 10-, or 15-mM glucose. **(B)** Glucagon secretion in human islets (n=5) treated with 3 mM R- or S- βHB for 72 h, at 3-, 6-, or 10 mM glucose. Each circle represents a technical replicate (islets from the same donor). **(C-E)** Insulin secretion in human islets treated with 3 mM R-βHB for 72 h at 3 mM glucose (C), 6 mM glucose (D), or 10 mM glucose (E) stratified by BMI. **(F-H)** Insulin secretion in human islets treated with 3 mM S-βHB for 72 h at 3 mM glucose (E), 6 mM glucose (F), or 10 mM glucose (G) stratified by BMI. Each colour corresponds to an individual donor. Filled circles correspond to donors with BMI>25, and hollow circles correspond to donors with BMI<25. The BMI of individual donors is listed to the right of each plot. Each point is the average of 3-4 technical replicates.

### Supplementary Figure 6

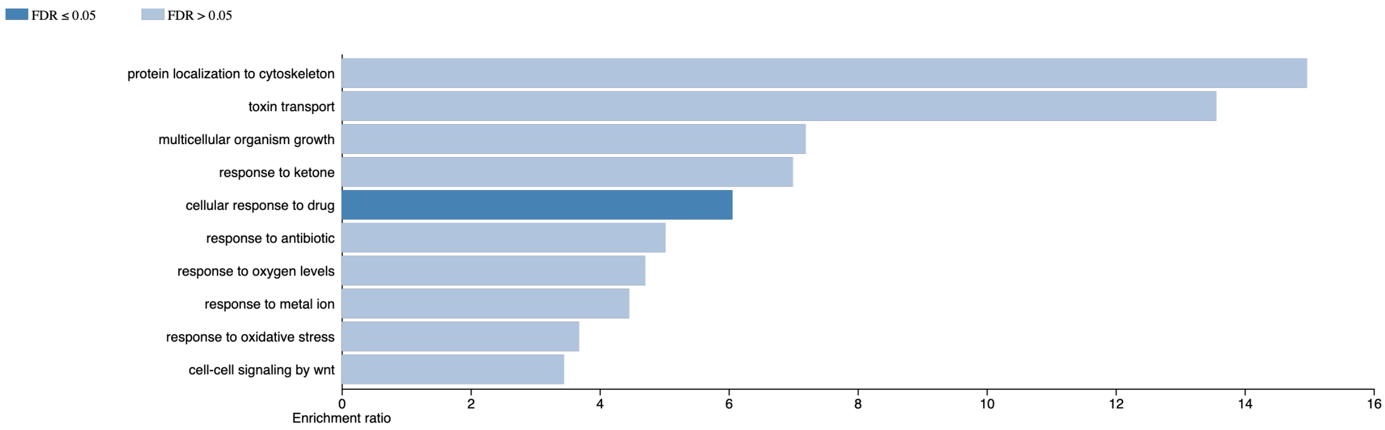


**Supplementary Figure 6. Proteomics pathway analysis.** WebGestalt Over-Representation Analysis (ORA) shows biological processes involving differentially abundant proteins in islets treated with R-βHB for 72 h ($p<0.05$). Enrichment ratio indicates the ratio of the number of proteins submitted to WebGestalt to the total number of proteins in that category on Kyoto Encyclopedia of Genes and Genomes (KEGG).

Supplementary Figure 7

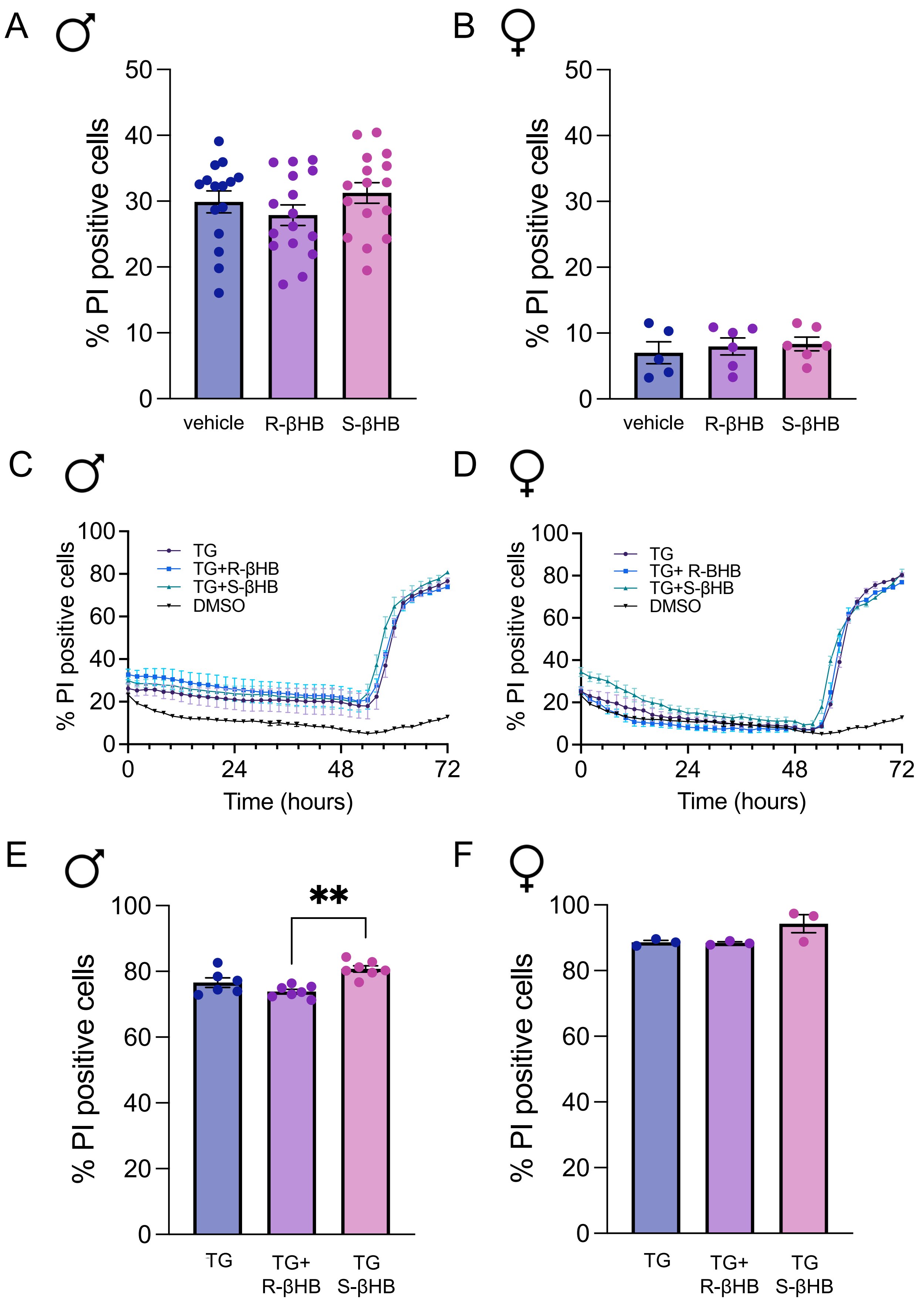

**Supplementary Figure 7. Effect of individual enantiomers of βHB on cell death in dispersed islet cells isolated from male and female mice.** Cell death as % PI positive cells was quantified at the final time point in dispersed islet cells treated with either enantiomer of βHB **(A-B)** Cell death as % PI positive cells was quantified at the final time point in dispersed islet cells from male mice (A) and female mice (B). **(C-D)** Cell death was assessed in dispersed islet cells treated with 10 μM TG ± R-βHB or S-βHB and separated by sex (n=2 male, n=1 female mice). **(E-F)** Cell death quantified as % PI positive cells at 72 h in dispersed islet cells treated with 10 μM TG ± R-βHB or S-βHB from male (E) and female (F) mice. Bars represent mean ± SEM, each circle represents a technical replicate. Statistical significance is indicated as asterisks above the corresponding bars. ** indicates $p<0.01$.
